# tangermeme: A toolkit for understanding *cis*-regulatory logic using deep learning models

**DOI:** 10.1101/2025.08.08.669296

**Authors:** Jacob Schreiber

## Abstract

Deep learning models have achieved state-of-the-art performance at predicting diverse genomic modalities, yet their promise for biological discovery lies in how they are used *after* demonstrating their predictive performance. Here, we describe the functionality of tangermeme, a highly optimized toolkit for “everything-but-the-model” when it comes to genomic deep learning, and demonstrate how tangermeme can be used to distill the learned *cis-* regulatory patterns from models into human-interpretable insights.

## Main

Machine learning models, such as hidden Markov models [1] and support vector machines [2, 3], have been important tools in genomics for decades [4]. However, the rapid adoption of deep learning has marked a fundamental shift in how machine learning is applied to genomics data [5, 6, 7, 8]. An important strength of deep learning models in genomics is their end-to-end nature, where manual feature engineering is replaced with training models directly on nucleotide sequence. Consequently, state-of-the-art models now exist that make predictions for transcription factor (TF) binding [9], histone modification [10], chromatin accessibility [11, 12, 13] and architecture [14, 15], transcription [16, 17], alternative splicing [18, 19], the binding of miRNAs [20], RNA degredation rates [21], including many of these forms of activity at single cell or spatial resolutions.

Although most attention has been paid to achieving the best predictive performance with these models, this is only the first step on the path to biological discovery [22]. After showing that these models are accurate, they can estimate the effect of non-coding variants and identify causal drivers [23, 24], paired with feature attribution methods to identify the nucleotides/motifs driving model predictions [25, 26], probed to identify global *cis-*regulatory rules [27, 28], or used to design sequences with desired characteristics [29, 30, 31]. Crucially, this downstream usage is usually agnostic to the details of model training; the process for estimating non-coding variant effect is largely unaffected by whether the model contains convolutions or transformers or by the choice of optimizer.

Despite this modularity, there does not yet exist an optimized repository for such methods that is generalizable across model types. Instead, bespoke analysis code is usually packaged alongside models when they are released. Some software packages, such as Captum [32] implement one or a small number of related algorithms. Other packages such as Selene [33], CREsted [34], EUGENe [35], and gReLU [36] focus on the training and fine-tuning of models, with some even including model zoos of pre-trained models. Although being able to train or adapt models to a novel research setting is clearly invaluable, their support for using these models *after* training is much more limited.

Accordingly, we introduce tangermeme, a software package which implements “everything-but-the-model” when it comes to genomic deep learning. This design choice is intentional, as the computational layers within models and their training strategies are much more diverse and rapidly evolving than how these models are subsequently used. Instead, tangermeme focuses on using trained models to drive genomics research (Fig 1A), including support for sequence manipulations, model operations, and efficient combinations of the two (e.g. variant effect prediction). These operations have built-in batching, support for alternative data types and devices, and work out-of-the-box on multi-input/output models. Each operation is low-level enough to be easily extended or integrated into research code bases.

**Figure 1.**
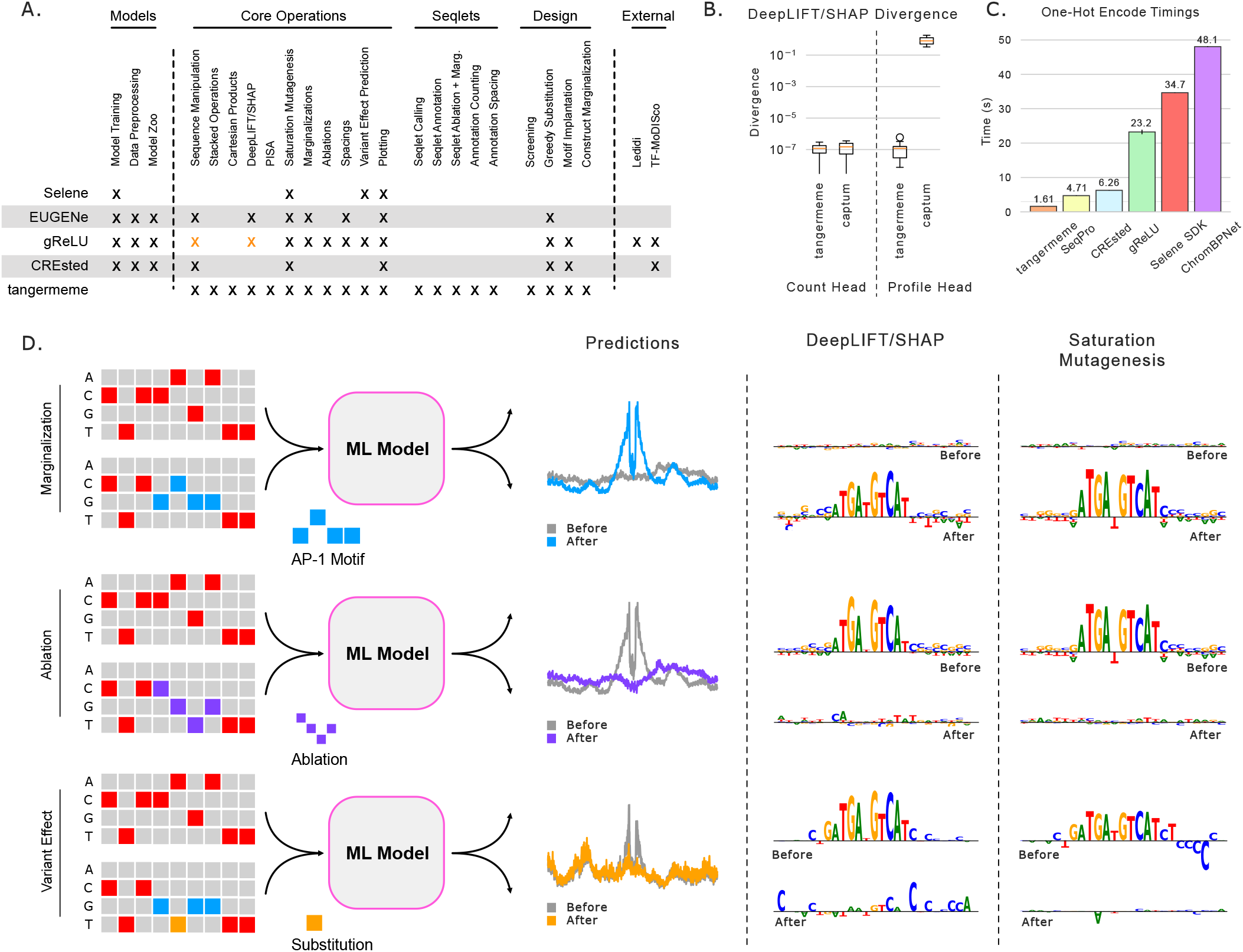
An overview of the tangermeme software suite. (A) A comparison of functionality. Orange X’s indicates functionality that uses tangermeme. (B) Covergence deltas from DeepLIFT/SHAP in two settings. (C) Timings for one-hot encoding hg38 chr1, averaged across ten runs. Standard errors are plotted but, except for gReLU, are too small to easily see. (D) A cartoon schematic and simulation results demontrating that sequence manipulations (the rows) can be separated by any model operation (the columns).

Each of tangermeme’s operations are designed to simplify the use of deep learning in genomics. For instance, most DeepLIFT/SHAP implementations require all background sequences in GPU memory simultaneously, making massive models difficult to interpret. Tangermeme’s built-in batching removes this requirement without compromising exactness, and lowprecision support further reduces the burden. Further, these implementations can silently fail, causing the convergence deltas to become quite large instead of near zero [37] (Fig 1B), whereas tangermeme will never do so. Finally, core operations in tangermeme are significantly faster than other software. It takes tangermeme less than two seconds to one-hot encode the entirety of hg38 chr1 (Fig 1C) — almost three times faster than the next fastest implementation. Although one-hot encoding is conceptually simple, being this fast enables on-the-fly batch generation and is reflective of the optimizations taken across tangermeme.

An important consequence of tangermeme’s design is that operations can be stacked. Typically, variant effect prediction implementations substitute a variant into a reference sequence and then make predictions. However, tangermeme does not require that this second step is prediction. By instead calculating attributions, one can gain insight into how a variant may disrupt *cis-*regulatory logic formed through several motifs (Fig 1D). By separating the sequence manipulation and model operation steps of common procedures, tangermeme opens up an understudied class of analyses that will be broadly useful to researchers.

In addition to implementing commonly used methods, tangermeme includes novel algorithms for automatically distilling learned *cis-*regulatory logic from trained deep learning models. These algorithms focus on the calling and usage of “seqlets”, which are contiguous spans of high-attribution characters from an algorithm such as DeepLIFT/SHAP, *in silico* saturation mutagenesis, PISA [38], or saliency [39] (Fig 2A). This seqlet caller is the first to call variable-length seqlets directly, and does so using a principled definition of a seqlet to provide a simple and robust algorithm (see Methods for details). After seqlets are called, tangermeme includes steps for annotating them using a motif database and then deriving statistics from these annotations, such as counts, pairwise presence, and spacing relationships.

**Figure 2.**
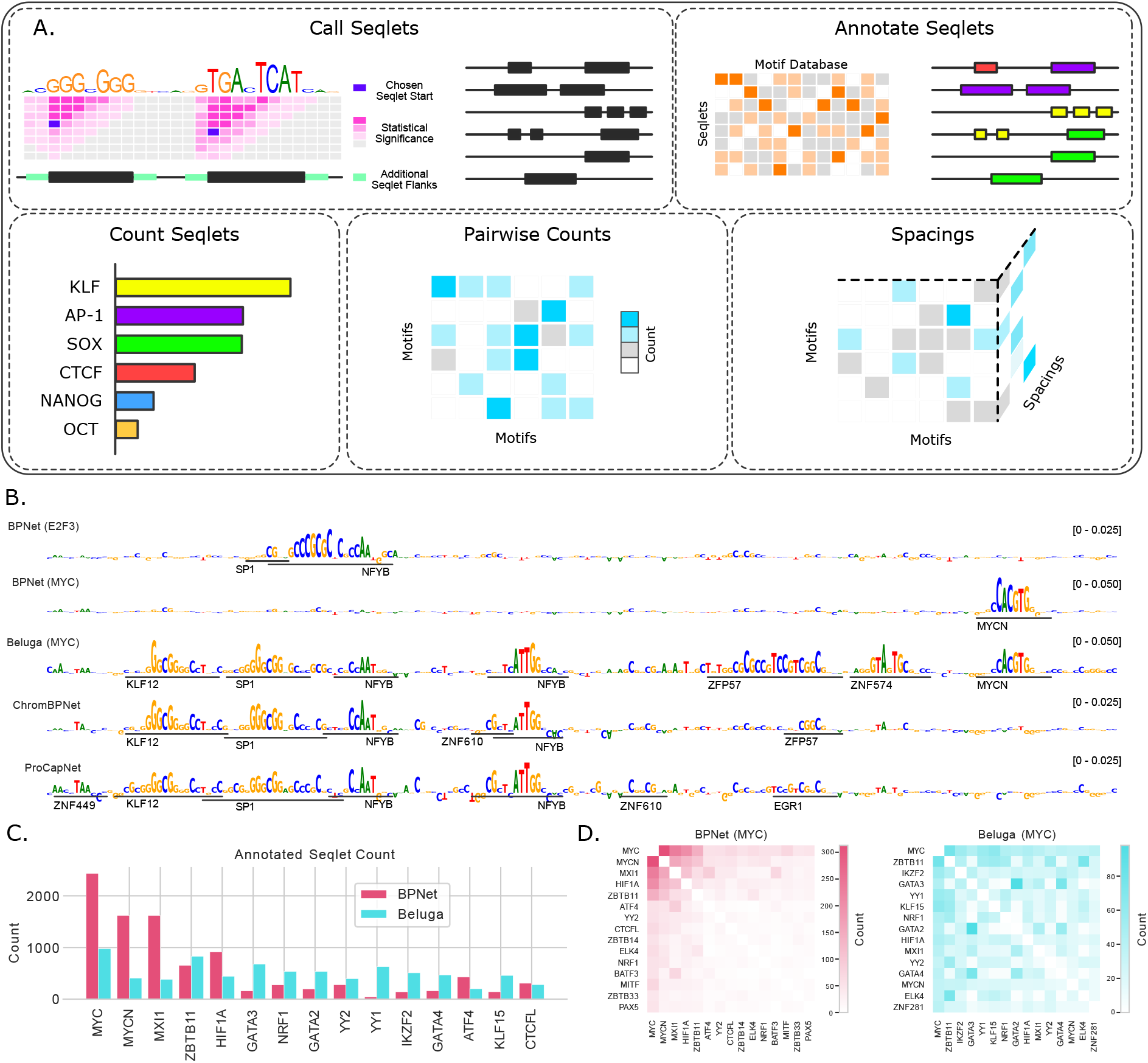
An automatic pipeline for the identification and usage of seqlets. (A) An overview of the implemented steps. (B) Example attributions at the PLD6 promoter along with seqlet calls (the bars underneath) and annotations. (C) Motif counts for BPNet and Beluga models predicting MYC binding. (D) Pairwise motif counts for the models in C showing the top 15 motifs for each.

As a demonstration of these combined capabilities, we used tangermeme to compare the learned logics of several models (see Methods for details). Attributions revealed that alternate sets of motifs drove predictions across different models at the promoter of PLD6 with, interestingly, different motifs driving MYC predictions from BPNet [9] and Beluga [24] (Fig 2B). By automatically calling and annotating seqlets using these new algorithms, we can see that BP-Net’s predictions are solely driven by a MYC motif whereas Beluga is driven by many motifs that also drive accessibility and transcription initiation. We can then more comprehensively describe the motifs learned by these models by counting the annotated seqlets across all MYC peaks (Fig 2C) and also through pairwise occurrences of these motifs (Fig 2D). Through this automatic procedure, we can immediately see that the learned logics for BPNet and Beluga are significantly different despite being ostensibly trained to predict the same phenomena, with BPNet focusing almost exclusively on MYC motifs and Beluga learning a broader lexicon of motifs and interactions also associated with chromatin accessibility.

Tangermeme is a software package for genomic deep learning that offers efficient implementations of core algorithms as well as novel algorithms for distilling learned *cis-*regulatory patterns into insights. Its operations are easy-to-use, well documented, and can be adapted or extended to research code in a variety of settings. We anticipate that tangermeme will be increasingly valuable as researchers continue to build state-of-the-art deep learning models in genomics, and that it will enable deeper analysis and biological exploration to be performed.

## Methods

### One-hot Encodings

We compared the speed of one-hot encoding the entirety of hg38 chr1 using several software packages. These included tangermeme (v0.5.1), SeqPro (v0.6.1), CREsted (v1.4.0), gReLU (v1.0.7), and Selene SDK (0.6.0). We also used the one-hot encoding function in ChromBPNet (v1.0.1), although we did not install the entire package due to its TensorFlow dependency.

Each package was timed solely on its respective one-hot encoding function. For several packages, including SeqPro and Selene SDK, there were steps involved with defining an alphabet that were not included in the timings. Each timing was performed on a single thread to avoid issues related to parallel processing that may have arisen.

### Models

In this work, we used five deep learning models, which are publicly available at https://zenodo.org/records/16751681. We trained the two BPNet models and are re-hosting the other three models for convenience.

We trained BPNet models to predict the binding of E2F3 and MYC using the default parameters in the bpnet-lite v0.9.0 command-line tool. BPNet models make predictions for basepair resolution profiles, which are the probability of an observed read mapping to each nucleotide in the output window of the model, and also for the log of the total number of read counts within the window. Respectively, these are the shape and the strength of the binding. These models were trained on ChIP-seq experiments hosted on the ENCODE Compendium [40] (ENCSR036QIR and ENCSR000EGJ for E2F3 and MYC respectively, both using the control track ENCSR000BLJ). In each case, loci for training included those in the peaks file associated with each experiment along with GC-matched negatives computed from those peaks files. During training, 2114 bp windows were extracted as input to the models that were centered on the peaks or chosen background regions *±* a randomly chosen jitter between -128 and 128. Half of the seen examples were reverse-complemented, meaning that both the nucleotide sequence and the basepair resolution profiles were flipped along both the length and feature axes. A fixed count loss weight of 100 was used.

The DeepSEA/Beluga model was downloaded from Kipoi [41] and modified to change the expected AGCT alphabet to be the more popular ACGT alphabet (http://kipoi.org/models/DeepSEA/beluga/). We considered only the output for MYC in K562 (target 614, 0-indexed)

The ChromBPNet and ProCapNet models were downloaded from the ENCODE Compendium (https://www.encodeproject.org/annotations/ENCSR467RSV/ and https://www.encodeproject.org/annotations/ENCSR740IPL/respectively). In both cases, we used models from only fold 0.

### Simulations

In Fig 1, we showed how tangermeme decouples sequence manipulatons from the subsequent model operations to enable a richer space of analyses. In these simulations we show predictions and attributions from the profile prediction output of a ChromBPNet model. For the marginalizations and ablations, we used the same set of 100 randomly generated DNA sequences that were 2,114bp long using nucleotide probabilities derived from the genome (28% A/T, 22% C/G). For the marginalizations, we performed the respective operation with ChromBPNet before and after substituting “ATGATGTCAT” into the middle of each of these sequences and display both, averaged across the sequences. For the ablations, we began by substituting that same subsequence into the sequences to generate the “before” sequences and then dinucleotide shuffled the window between 1030bp and 1075bp to get the “after” sequences. When running DeepLIFT/SHAP we used only one background shuffle per sequence because we are already averaging across the 100 randomly generated sequences. When running saturation mutagenesis, we perform a windowed version that focuses solely on the 1040-1075bp window for speed without losing exactness, which surrounds where the motif is being substituted into. For the variant effect predictions, we change the nucleotide at position 1053 from a G to an A.

### Seqlet Calling

Tangermeme includes a novel seqlet calling algorithm that uses a recursive application of a basic statistical test to call seqlets from attribution scores. This recursive application makes it the first seqlet calling algorithm to be able to call variable length seqlets directly. Briefly, a seqlet is a contiguous span of nucleotides with high attribution values. Our seqlet caller further defines a seqlet as a span with a statistically significant attribution sum compared to a background distribution, where *all subspans* within the seqlet would also be called seqlets according to this statistical test. This property does not need to hold for subspans below a certain length (by default 4) to prevent noise in the attributions from disrupting a seqlet call and to intentionally allow seqlets with uninformative characters within them.

This simple definition of a seqlet leads to a principled calling algorithm with only a few parameters: the p-value threshold, the minimum and maximum seqlet lengths, optional additional flanks to add to the edges for robustness in downstream processing, and the number of bins to use when converting the contiguous signal to a discrete one. Of these, the p-value threshold is likely the most important to tune, although the minimum seqlet length may also matter depending on how noisy the attributions are. Changes to the maximum seqlet length do not affect seqlets that are smaller than the current maximum, and the number of bins is broadly robust across a range of values and mostly influences computatonal time and memory.

The algorithm proceeds as follows. We begin with attribution values *X* ∈ ℝ^*n,l*^ where *n* is the number of examples and *l* is the length of these examples. In principle, *l* does not need to be constant across examples but in our current implementation it is assumed to be fixed. These attribution values are then binned by min-max scaling them using the global min and max values, multiplying by the number of bins *b*, and taking the floor of the resulting value.

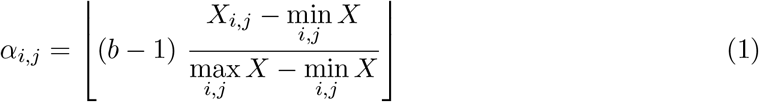

These integerized scores are then used to create a frequency histogram *f* ∈ ℝ^*b*^:

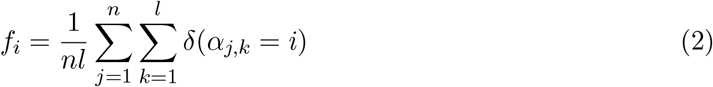

where *δ* is the Kronecker delta function which is 1 when the term is true and 0 otherwise. This is similar to the initial steps in the procedure in Tomtom [42] for calculating background distributions.

This frequency histogram is then used to calculate null distributions for each seqlet length. These null distributions essentially answer the question “what attribution sum would I get by randomly choosing this number of (integerized versions of the) attribution values from the data?” without considering whether the values are contiguous. Each null distribution encodes a PDF over integerized scores, with the span of these scores increasing with length. Specifically, each null distribution spans between 0 and *tb* with *t* being the seqlet length. For simplicity, we will represent all of these PDFs in a single sparse matrix *A* ∈ ℝ^*T,T b*^ where *T* is the maximum seqlet length and *A*_1,0:*b*_ is set to be equal to *f* and the rest of *A* is filled in by

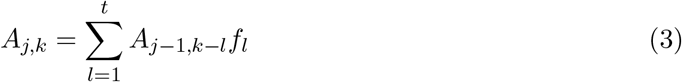

Although this procedure is iterative in nature, it is not the inspiration for the use of “recursive” in the name of the seqlet caller.

Each of these null distributions is currently a PDF. Because we will need to quickly calculate p-values, we convert these PDFs into 1 - CDFs from which determining p-values is a simple indexing operation.

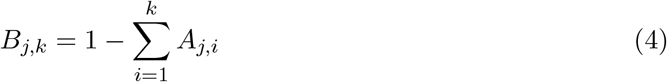

Given these 1 - CDFs, we can then convert our original attributions *X* into p-values by binning them using the same procedure as before and indexing into these *B* distributions. This yields *P* ∈ ℝ^*n,l*^ comprised of p-values for each position and each potential seqlet span length. Seqlets are then called from *P* based on the definition that seqlets are entries in *P* whose value falls below the threshold, and also where all subspans (longer than the minimum seqlet span) within the seqlet also fall below the threshold. For efficiency, this is done by setting the first rows between 0 and `min_seqlet_len` to 0 and then setting each entry *P*_*i,j*_ to 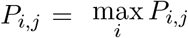. Now, by indexing into each element, we can know whether all spans beginning at that position pass the p-value threshold. We can do this efficiently by proceeding from *i* = 0 to *i* = `max_seqlet_len` and setting *P*_*i,j*_ = max(*P*_*i,j*_, *P*_*i*−1,*j*_). If any spans are not statistically significant, this procedure will block any longer spans from being considered. To consider *all* subspans and not just those starting at position *j*, we then proceed diagonally from right to left, setting *P*_*i,j*_ = max(*P*_*i,j*_, *P*_*i*−*i,j*+1_). Together, these two steps mean that when the value at *P*_*i,j*_ is below the threshold, all subspans within the width would be called as statistically significant. Accordingly, we can proceed to call seqlets by iterating from the maximum seqlet length to the minimum seqlet length, checking whether the minimum p-value for that row is below the threshold, and when identifying a seqlet setting all columns within the width of the seqlet to 1 to prevent multiple seqlets from being called in the same position. This procedure means that the *longest* seqlet will be called for a given set of positions, not necessarily the most statistically significant.

### Seqlet Annotation

Once seqlets have been called using the recursive seqlet caller they are then annotated with the motif they most resemble. This is done using the `annotate_seqlets` command. This annotation procedure involves using tomtom-lite [43] to map the nucleotide sequence within the seqlet to a provided motif database. In these evaluations, the motif database was the slice of JASPAR2024 [44] database derived from ChIP-seq experiments performed in *Homo sapiens*. Seqlets were annotated with the motif with the lowest p-value.

### Seqlet Counting

After seqlets have been called and annotated, basic statistics can be extracted from the presence of these annotated seqlets across sequences. We can count the number of times each annotation appears in each sequence to get a *C* ∈ ℤ^*n,m*^ matrix where *n* is the number of sequences and *m* is the number of motifs in the motif database. This matrix can then be summed across the first dimension to count the number of times each motif is present, or across the second dimension to get the number of seqlets called per sequence. We show the first of these statistics for the BPNet (MYC) and Beluga (MYC) models in Fig 2C.

Going further, we can also count the number of times each pair of motifs appear within the same sequence and get *P* ∈ ℤ^*n,m,m*^. There are several modes this procedure can be run in. First, it can be run asymetrically or symetrically. When run symetrically, the presence of motifs A and B are counted in the row associated with A and also in the row associated with B, yielding a matrix that is symmetric. When run asymetrically, motifs are counted according to their order in the sequence from the start to the end, not accounting for the strandedness of the motif. This can be useful when the ordering of motifs is important, such as for transcription initiation, though one must manually ensure that the strandedness is correct. The default is running symmetrically. The second mode involves whether motif hits are unique or not. For example, when the motifs within a sequence are “A A A B”, whether we record three “A B” pairs or just a single one. This can be particularly important when dealing with arrays of motifs, such as KLF, because the count of those array elements may end up being significantly higher than others.

Finally, we can also consider the spacing between motifs. Here, we create a count of each pair of motifs at each possible spacing up until a user-defined maximum. All spacings after that maximum are ignored. Similar to the pairwise motif counts, this matrix can be symmetric or not. This matrix will likely be extremely sparse as many motifs do not co-occur at all, and even those that do co-occur are unlikely to do so frequently enough to fill in many potential spacings.

## Code Availability

tangermeme is free and open source software available under the MIT license at https://github.com/jmschrei/tangermeme. The latest version can be installed via pip using `pip install tangermeme`. Documentation and tutorials are available at https://tangermeme.readthedocs.io/en/latest/, including tutorials that produce the figures shown in this work.

## Acknowledgements

We thank Nikolaus Mandlburger, Alan Murphy, Seppe de Winter, Bo Hansen, and Adam He for their contributions to the code. We also thank Niklas Kempynck, Danila Bredikhin, Kelly Cochran, Habib Daneshpajouh, Selin Jessa, Damla Ö vek Baydar, and the rest of the tangermeme community for their feedback. Research at the Institute of Molecular Pathology (IMP) is supported by Boehringer Ingelheim GmbH and the Austrian Research Promotion Agency (FFG, FO999902549). For the purpose of Open Access, the authors have applied a CC BY public copyright license to any Author Accepted Manuscript (AAM) version arising from this submission.

## References

[1] Stephen Winters-Hilt. Hidden markov model variants and their application. BMC Bioinformatics, 7 Suppl 2(S2):S14, September 2006.

[2] Shujun Huang, Nianguang Cai, Pedro Penzuti Pacheco, Shavira Narrandes, Yang Wang, and Wayne Xu. Applications of support vector machine (SVM) learning in cancer genomics. Cancer Genomics Proteomics, 15(1):41–51, January 2018.

[3] William S Noble. What is a support vector machine? Nat. Biotechnol., 24(12):1565–1567, December 2006.

[4] Maxwell W Libbrecht and William Stafford Noble. Machine learning applications in genetics and genomics. Nat. Rev. Genet., 16(6):321–332, June 2015.

[5] Tristan Bepler and Bonnie Berger. Learning the protein language: Evolution, structure, and function. Cell Syst., 12(6):654–669.e3, June 2021.

[6] Britta Velten and Oliver Stegle. Principles and challenges of modeling temporal and spatial omics data. Nat. Methods, September 2023.

[7] Gökcen Eraslan, Žiga Avsec, Julien Gagneur, and Fabian J Theis. Deep learning: new computational modelling techniques for genomics. Nat. Rev. Genet., 20(7):389–403, July 2019.

[8] Xuehai Hu, Alisdair R Fernie, and Jianbing Yan. Deep learning in regulatory genomics: from identification to design. Current Opinion in Biotechnology, 79:102887, February 2023.

[9] Žiga Avsec, Melanie Weilert, Avanti Shrikumar, Sabrina Krueger, Amr Alexandari, Khyati Dalal, Robin Fropf, Charles McAnany, Julien Gagneur, Anshul Kundaje, and Julia Zeitlinger. Base-resolution models of transcription-factor binding reveal soft motif syntax. Nat. Genet., 53(3):354–366, March 2021.

[10] Qijin Yin, Mengmeng Wu, Qiao Liu, Hairong Lv, and Rui Jiang. DeepHistone: a deep learning approach to predicting histone modifications. BMC Genomics, 20(Suppl 2):193, April 2019.

[11] David R Kelley, Jasper Snoek, and John L Rinn. Basset: learning the regulatory code of the accessible genome with deep convolutional neural networks. Genome Res., 26(7):990–999, July 2016.

[12] Anusri Pampari, Anna Shcherbina, Evgeny Z Kvon, Michael Kosicki, Surag Nair, Soumya Kundu, Arwa S Kathiria, Viviana I Risca, Kristiina Kuningas, Kaur Alasoo, William James Greenleaf, Len A Pennacchio, and Anshul Kundaje. ChromBPNet: bias factorized, base-resolution deep learning models of chromatin accessibility reveal cisregulatory sequence syntax, transcription factor footprints and regulatory variants. bioRxivorg, page 2024.12.25.630221, January 2025.

[13] Yan Hu, Max A Horlbeck, Ruochi Zhang, Sai Ma, Rojesh Shrestha, Vinay K Kartha, Fabiana M Duarte, Conrad Hock, Rachel E Savage, Ajay Labade, Heidi Kletzien, Alia Meliki, Andrew Castillo, Neva C Durand, Eugenio Mattei, Lauren J Anderson, Tristan Tay, Andrew S Earl, Noam Shoresh, Charles B Epstein, Amy J Wagers, and Jason D Buenrostro. Multiscale footprints reveal the organization of cis-regulatory elements. Nature, 638(8051):779–786, February 2025.

[14] Jian Zhou. Sequence-based modeling of three-dimensional genome architecture from kilo-base to chromosome scale. Nat. Genet., 54(5):725–734, May 2022.

[15] Geoff Fudenberg, David R Kelley, and Katherine S Pollard. Predicting 3D genome folding from DNA sequence with akita. Nat. Methods, 17(11):1111–1117, November 2020.

[16] Bernardo P de Almeida, Franziska Reiter, Michaela Pagani, and Alexander Stark. Deep-STARR predicts enhancer activity from DNA sequence and enables the de novo design of synthetic enhancers. Nat. Genet., 54(5):613–624, May 2022.

[17] Kelly Cochran, Melody Yin, Anika Mantripragada, Jacob Schreiber, Georgi K Marinov, and Anshul Kundaje. Dissecting the cis-regulatory syntax of transcription initiation with deep learning. bioRxivorg, page 2024.05.28.596138, May 2024.

[18] Kishore Jaganathan, Sofia Kyriazopoulou Panagiotopoulou, Jeremy F McRae, Siavash Fazel Darbandi, David Knowles, Yang I Li, Jack A Kosmicki, Juan Arbelaez, Wenwu Cui, Grace B Schwartz, Eric D Chow, Efstathios Kanterakis, Hong Gao, Amirali Kia, Serafim Batzoglou, Stephan J Sanders, and Kyle Kai-How Farh. Predicting splicing from primary sequence with deep learning. Cell, 176(3):535–548.e24, January 2019.

[19] Tony Zeng and Yang I Li. Predicting RNA splicing from DNA sequence using pangolin. Genome Biol., 23(1):103, April 2022.

[20] Bhargav Kanuparthi, Sara E Pour, Scott D Findlay, Omar Wagih, Jahir M Gutierrez, Rory Gao, Jeff Wintersinger, Junru Lin, Martino Gabra, Emma Bohn, Tammy Lau, Chris Cole, Andrew Jung, Albi Celaj, Fraser Soares, Rachel Gray, Brandon Vaz, Kate Delfosse, Varun Lodaya, Sakshi Bhargava, Diane Ly, Farhan Yusuf, Ken Kron, Greg Hoffman, Shreshth Gandhi, and Brendan J Frey. Sequence based prediction of cell type specific microRNA binding and mRNA degradation for therapeutic discovery. bioRxiv, page 2025.05.15.654105, May 2025.

[21] Vikram Agarwal and David R Kelley. The genetic and biochemical determinants of mRNA degradation rates in mammals. Genome Biol., 23(1):245, November 2022.

[22] Gherman Novakovsky, Nick Dexter, Maxwell W Libbrecht, Wyeth W Wasserman, and Sara Mostafavi. Obtaining genetics insights from deep learning via explainable artificial intelligence. Nat. Rev. Genet., 24(2):125–137, February 2023.

[23] Xiaoyu Wang, Fuyi Li, Yiwen Zhang, Seiya Imoto, Hsin-Hui Shen, Shanshan Li, Yuming Guo, Jian Yang, and Jiangning Song. Deep learning approaches for non-coding genetic variant effect prediction: current progress and future prospects. Brief. Bioinform., 25(5):bbae446, July 2024.

[24] Jian Zhou and Olga G Troyanskaya. Predicting effects of noncoding variants with deep learning-based sequence model. Nat. Methods, 12(10):931–934, October 2015.

[25] Ferran Muiños, Francisco Martínez-Jiménez, Oriol Pich, Abel Gonzalez-Perez, and Nuria Lopez-Bigas. In silico saturation mutagenesis of cancer genes. Nature, 596(7872):428–432, August 2021.

[26] Avanti Shrikumar, Peyton Greenside, and Anshul Kundaje. Learning important features through propagating activation differences. arXiv [cs.CV], April 2017.

[27] Peter K Koo, Antonio Majdandzic, Matthew Ploenzke, Praveen Anand, and Steffan B Paul. Global importance analysis: An interpretability method to quantify importance of genomic features in deep neural networks. PLoS Comput. Biol., 17(5):e1008925, May 2021.

[28] Shushan Toneyan and Peter K Koo. Interpreting cis-regulatory interactions from largescale deep neural networks. Nat Genet, pages 1–11, September 2024.

[29] Jacob Schreiber, Franziska Katharina Lorbeer, Monika Heinzl, Yang Lu, Alexander Stark, and William Stafford Noble. Programmatic design and editing of cis-regulatory elements. bioRxiv, page 2025.04. 22.650035, April 2025.

[30] Johannes Linder and Georg Seelig. Fast activation maximization for molecular sequence design. BMC Bioinformatics, 22(1):510, October 2021.

[31] Seppe De Winter, Vasileios Konstantakos, and Stein Aerts. Modelling and design of transcriptional enhancers. Nat Rev Bioeng, pages 1–16, February 2025.

[32] Narine Kokhlikyan, Vivek Miglani, Miguel Martin, Edward Wang, Bilal Alsallakh, Jonathan Reynolds, Alexander Melnikov, Natalia Kliushkina, Carlos Araya, Siqi Yan, and Orion Reblitz-Richardson. Captum: A unified and generic model interpretability library for PyTorch. arXiv [cs.LG], September 2020.

[33] Kathleen M Chen, Evan M Cofer, Jian Zhou, and Olga G Troyanskaya. Selene: a PyTorch-based deep learning library for sequence data. Nat. Methods, 16(4):315–318, April 2019.

[34] Niklas Kempynck, Lukas Mahieu, Eren Can Ekşi, Vasilis Konstantakos, Cas Blaauw, Seppe De Winter, Gert Hulselmans, Ibrahim Taskiran, and Stein Aerts. CREsted: Cis regulatory element sequence training, explanation, and design, 2024.

[35] Adam Klie, David Laub, James V Talwar, Hayden Stites, Tobias Jores, Joe J Solvason, Emma K Farley, and Hannah Carter. Predictive analyses of regulatory sequences with EUGENe. Nat. Comput. Sci., 3(11):946–956, November 2023.

[36] Avantika Lal, Laura Gunsalus, Surag Nair, Tommaso Biancalani, and Gokcen Eraslan. gReLU: A comprehensive framework for DNA sequence modeling and design. bioRxiv, page 2024.09.18.613778, September 2024.

[37] Jacob Schreiber. Attribution trickiness and DeepLiftShap implementations, 2024.

[38] Charles E McAnany, Melanie Weilert, Grishma Mehta, Fahad Kamulegeya, Jennifer M Gardner, Anshul Kundaje, and Julia Zeitlinger. PISA: a versatile interpretation tool for visualizing cis-regulatory rules in genomic data. bioRxiv, page 2025.04.07.647613, April 2025.

[39] Antonio Majdandzic, Chandana Rajesh, and Peter K Koo. Correcting gradient-based interpretations of deep neural networks for genomics. Genome Biol., 24(1):109, May 2023.

[40] ENCODE Project Consortium, Jill E Moore, Michael J Purcaro, Henry E Pratt, Charles B Epstein, Noam Shoresh, Jessika Adrian, Trupti Kawli, Carrie A Davis, Alexander Dobin, Rajinder Kaul, Jessica Halow, Eric L Van Nostrand, Peter Freese, David U Gorkin, Yin Shen, Yupeng He, Mark Mackiewicz, Florencia Pauli-Behn, Brian A Williams, Ali Mortazavi, Cheryl A Keller, Xiao-Ou Zhang, Shaimae I Elhajjajy, Jack Huey, Diane E Dickel, Valentina Snetkova, Xintao Wei, Xiaofeng Wang, Juan Carlos Rivera-Mulia, Joel Rozowsky, Jing Zhang, Surya B Chhetri, Jialing Zhang, Alec Victorsen, Kevin P White, Axel Visel, Gene W Yeo, Christopher B Burge, Eric Lécuyer, David M Gilbert, Job Dekker, John Rinn, Eric M Mendenhall, Joseph R Ecker, Manolis Kellis, Robert J Klein, William S Noble, Anshul Kundaje, Roderic Guigó, Peggy J Farnham, J Michael Cherry, Richard M Myers, Bing Ren, Brenton R Graveley, Mark B Gerstein, Len A Pennacchio, Michael P Snyder, Bradley E Bernstein, Barbara Wold, Ross C Hardison, Thomas R Gingeras, John A Stamatoyannopoulos, and Zhiping Weng. Expanded encyclopaedias of DNA elements in the human and mouse genomes. Nature, 583(7818):699–710, July 2020.

[41] Žiga Avsec, Roman Kreuzhuber, Johnny Israeli, Nancy Xu, Jun Cheng, Avanti Shrikumar, Abhimanyu Banerjee, Daniel S Kim, Thorsten Beier, Lara Urban, Anshul Kundaje, Oliver Stegle, and Julien Gagneur. The kipoi repository accelerates community exchange and reuse of predictive models for genomics. Nat. Biotechnol., 37(6):592–600, June 2019.

[42] Shobhit Gupta, John A Stamatoyannopoulos, Timothy L Bailey, and William Stafford Noble. Quantifying similarity between motifs. Genome Biol., 8(2):R24, February 2007.

[43] Jacob Schreiber. Tomtom-lite: Accelerating tomtom enables large-scale and real-time motif similarity scoring. bioRxiv, page 2025.05.27.656386, May 2025.

[44] Ieva Rauluseviciute, Rafael Riudavets-Puig, Romain Blanc-Mathieu, Jaime A Castro-Mondragon, Katalin Ferenc, Vipin Kumar, Roza Berhanu Lemma, Jérémy Lucas, Jeanne Chèneby, Damir Baranasic, Aziz Khan, Oriol Fornes, Sveinung Gundersen, Morten Johansen, Eivind Hovig, Boris Lenhard, Albin Sandelin, Wyeth W Wasserman, François Parcy, and Anthony Mathelier. JASPAR 2024: 20th anniversary of the open-access database of transcription factor binding profiles. Nucleic Acids Res., 52(D1):D174–D182, January 2024.

